# An NMR fingerprint matching approach for the identification and structural re-evaluation of *Pseudomonas* lipopeptides

**DOI:** 10.1101/2022.01.07.475420

**Authors:** Vic De Roo, Yentl Verleysen, Benjámin Kovács, De Vleeschouwer Matthias, Léa Girard, Monica Höfte, René De Mot, Annemieke Madder, Niels Geudens, José C. Martins

## Abstract

Cyclic lipopeptides (CLiPs) are secondary metabolites secreted by a range of bacterial phyla. CLiPs from *Pseudomonas* in particular display diverse structural variations in terms of the number of amino acid residues, macrocycle size, amino acid identity and stereochemistry (e.g. D- vs. L-amino acids). Reports detailing the discovery of novel or already characterized CLiPs from new sources appear regularly in literature. Increasingly however, the lack of detailed characterization threatens to cause considerable confusion, especially if configurational heterogeneity is present for one or more amino acids. Using *Pseudomonas* CLiPs from the Bananamide, Orfamide and Xantholysin groups as test cases, we demonstrate and validate that ^1^H and ^13^C NMR chemical shifts of CLiPs are sufficiently sensitive to differentiate between possible diastereomers of a particular sequence even when they only differ in a single D/L configuration. Rapid screening, involving simple comparison of the NMR fingerprint of a newly isolated CLiP with that of a reference CLiP of known stereochemistry, can then be applied to resolve dead-ends in configurational characterization and avoid the much more cumbersome chemical characterization protocols. Even when the stereochemistry of a particular reference CLiP remains to be established, NMR fingerprinting still allows verifying whether a CLiP from a novel source is already present in the reference collection, thus contributing to dereplication. To benefit research involving CLiPs, we have made a publicly available ‘knowledge base’ at https://www.rhizoclip.be, where we present an overview of published NMR fingerprint data of characterized CLiPs, together with literature data on the originally determined structures.

**Significance Statement:** *Pseudomonas* CLiPs, are ubiquitous specialized metabolites, impacting the producer’s lifestyle and interactions with the (a)biotic environment. Consequently, they generate interest for agricultural and clinical applications. Establishing structure-activity relationships as premise to their development is hindered because full structural characterization including stereochemistry requires labor-intensive analyses, without guarantee for success. Moreover, increasing use of superficial comparison with previously characterized CLiPs introduces or propagates erroneous attributions, clouding further scientific progress. We provide a generally applicable characterization methodology for structural comparison of newly isolated CLiPs to reference compounds with (un)known stereochemistry based on NMR fingerprints. The reference compound database available for the wide scientific community promises to facilitate structural assessment and dereplication of newly isolated CLiPs, and to support genome mining for novel CLiPs.

## Introduction

Pseudomonads represent a diverse and ubiquitous bacterial genus that evolves in a wide range of ecological niches (1, 2). To support their lifestyle and interactions with other organisms, they produce a variety of complex secondary metabolites including cyclic lipodepsipeptides or ‘CLiPs’ generated by non-ribosomal peptide synthetases (NRPSs) (3). Biological effects associated with these biosurfactant peptides include antimicrobial activity, control of bacterial motility and biofilm formation and phytotoxicity, among others (4, 5). They are also involved in the complex interplay between bacteria and plants at the level of the plant rhizosphere. This generates plant-growth promoting activity and protection against pathogens by direct antagonism or induced resistance (6, 7). Other reports have detailed anti-insecticidal activity (8, 9). As a result, this diverse array of functions of *Pseudomonas* CLiPs has sparked interest in their potential development in plant biocontrol applications and as biopesticides (10, 11). In turn, this also stimulates screening efforts for their development as novel antimicrobials (4, 5). Indeed, the current dearth of novel compounds to fight multi-drug resistant bacteria or fungal infections combined with an increased focus on the development of narrow-spectrum or even species-specific antibiotics has renewed efforts towards mining bacteria for novel pharmaceutical leads (12). These have included CLiPs, (13, 14) not in the least motivated by the successful introduction of daptomycin into clinical settings. In addition, several reports have highlighted anti-carcinogenic activities of *Pseudomonas* CLiPs, further illustrating their biomedical application potential (15-17). Therefore, both from a fundamental and application perspective, there is a pressing need to uncover the mode of action resulting in these biological functions, starting with the way these are related to the underlying chemical structures (5, 18, 19). Given that structure elucidation of CLiPs remains far from trivial, the development of reliable and straightforward approaches remains key. Moreover, such approaches should allow comparing different CLiPs to unequivocally establish their structural identity.

With well over 100 distinct chemical structures reported to date, *Pseudomonas* CLiPs invariably consist of an oligopeptide sequence of 8 to 25 residues, which is partly cyclized into a depsipeptide through ester bond formation of the C-terminus with a preceding side-chain hydroxyl group (4, 5). An acyl chain of varying length and constitution caps the N-terminus of the oligopeptide sequence. The incorporation of non-proteinogenic amino acids, with a majority of residues displaying D-configuration through the action of epimerization domains in the non-ribosomal assembly line, generates additional structural diversity and further magnifies the structure elucidation challenge (20). The availability of rapid and affordable genome sequencing has stimulated the development of bioinformatic tools that allow to predict the CLiP structure produced by a particular NRPS by mapping its biosynthetic gene cluster (BGC) (21). The primary sequence is derived from analysis of the adenylation A-domains of the NRPS. Although separate epimerization domains are present in the siderophore synthetases of *Pseudomonas*, such activity is not found in their CLiP biosynthetic enzymes. Instead, condensation domains with intrinsic epimerization capacity (C/E) mimic the activity of a ^D^C_L_ domain by converting the configuration of the C-terminal residue in the intermediate from L to D before extending the peptide (22). Tentative attribution of D/L configuration can therefore be inferred from analysis of the condensation domains involved. For *Pseudomonas* CLiPs in particular, the dual condensation/epimerization C/E domains responsible for configurational inversion of the preceding L-amino acid in the growing peptide chain can be identified using such tools, however it turns out that the epimerization functionality can be inactive for reasons that remain unclear (23, 24). Consequently, as further demonstrated in this work, these predictions do not yet achieve the level of confidence necessary to eliminate the need for the more elaborate and labor-intensive chemical analysis methods. Furthermore, this genome-based approach does not provide information about the residues involved in the depsi bond, nor can the nature of the acyl chain be predicted. Thus, while analysis of the biosynthetic gene cluster coding for the NRPS represents a complementary asset during the chemical structure elucidation process (Figure S1), it cannot replace it.

Combined application of mass spectrometric (MS) and nuclear magnetic resonance (NMR) methods allows, in principle, to establish the composition of the oligopeptide sequence and the nature of the acyl chain as well as the cyclisation site but does not provide access to stereochemistry. For this, chemical derivatization methods combined with chromatographic separation, such as Marfey’s analysis, are typically employed to derive the number and configuration of individual amino acids (Figure S2)(25-27). However, the total hydrolysis of the lipopeptide into its constituent amino acids required in the process also causes the loss of information regarding their original position in the sequence. Consequently, positional ambiguity will arise when a particular amino acid with multiple occurrences in the sequence is configurationally heterogenous i.e., both the D- and L-configuration occur (26, 28). As a result, the elucidation of stereochemistry remains incomplete, such as is the case for the *Pseudomonas* CLiPs MDN-0066 (17), PPZPM-1a (29) and lokisin (30), among others. While workarounds exist, they are tedious and not error-proof (24, 26). In other cases, configurational analysis is omitted and configurational attribution is claimed by relying on either the conservation of the known pattern of D/L-configurations along a sequence (31, 32), or solely on genomic prediction (33-35). However, the latter ignores the problem of inactive epimerization domains (*vide supra*) while the former disregards the natural occurrence of diastereoisomeric lipopeptides having identical constitution but differing in configuration at one (or more) position(s). This is illustrated by, for example, the *Pseudomonas* CLiP pairs viscosin/WLIP, viscosinamide/pseudodesmin A, syringostatin/syringomycin and fengycin/plipastatin. Thus, new approaches that allow to move the structure elucidation beyond configurational dead-ends remain in high demand.

Simultaneously, the increased screening of *Pseudomonas* strains for CLiP production also calls for the introduction of dereplication approaches that confidently distinguish a known versus a novel CLiP structure early on, this to avoid the costly re-elucidation of previously described compounds (36). In line with this, multiple reports show a growing tendency to characterize known CLiPs from novel sources solely through comparison of their high-resolution mass to those of previously characterized ones (31, 32, 37-42). However, these approaches overlook the possibility of isobaric lipopeptides, which occur more commonly than is perhaps anticipated. For instance, several members of the Viscosin group of CLiPs, such as viscosin and massetolide F, have a distinct molecular topology yet the same molecular formula, a situation that also occurs for milkisin and tensin, both members of the Amphisin group. Even lipopeptides from distinct CLiP groups can be isobaric, such as strikingly demonstrated by gacamide/cocoyamide and putisolvin II (43-45). Indeed, these share the same C_66_H_115_N_13_O_19_ molecular formula yet differ in the total number of amino acids (12 vs. 11), the number of amino acids involved in the respective macrocycles (5 vs. 4) and the constitution of the acyl chain (3R-OH C10:0 vs. C6:0).

Previously, we noted that the combined ^1^H and ^13^C NMR chemical shift fingerprint obtained from ^1^H-{^13^C} HSQC spectra is sufficiently unique to differentiate diastereomers of a particular CLiP even when they differ in only a single D/L configuration (46). More so, it is sensitive to the stereochemistry of the 3-hydroxy moiety of the fatty acid tail as well (47). By integrating NMR spectral fingerprinting and total synthesis into the existing bioinformatic and chemical analyses workflows commonly used for structure elucidation we here show, for three representative cases, that configurational analysis dead-ends can be removed. First, by matching the ^1^H-{^13^C} HSQC spectral fingerprint of newly isolated *Pseudomonas* CLiPs with that of CLiPs of known stereochemistry obtained from total synthesis we demonstrated the strength of our approach on the CLiP produced by *Pseudomonas azadiae* SWRI103^T^. Next, the general applicability is shown by settling the stereochemistry of orfamide B from *Pseudomonas aestus* CMR5c and xantholysin A from *Pseudomonas mosselii* BW11M1, representative members from two other distinct *Pseudomonas* CLiP groups. In the process, we revise the stereochemistry of several orfamide homologues, including orfamide A from *P. protegens* Pf-5, the original prototype CLiP defining the Orfamide group (48). Next, we proceed to illustrate how NMR fingerprinting constitutes a potent tool for CLiP dereplication purposes by resolving configurational dead ends reported in literature for MDN-0066 (17), orfamides and xantholysin produced by various bacterial strains, without the need for additional synthesis. Based on this, we advocate the adoption of NMR based fingerprinting by the CLiP research community and initiate establishing a knowledge database, which will be further developed towards this purpose.

## Results

### 1. Structure elucidation of a newly isolated Bananamide group member from *P. azadiae* SWRI103

#### 1.1. Isolation, bioinformatics and initial chemical structure characterization

*P. azadiae* SWRI103 was originally collected from the rhizosphere of wheat in Iran, as part of a campaign to assess the plant growth stimulating potential of fluorescent pseudomonads (49). Initial PCR screening targeting initiation and termination domains indicated the presence of a lipopeptide-specific NRPS system (50). More recently, whole genome sequencing revealed a biosynthetic gene cluster (BGC) (51) with similarities to that of several Bananamide (8:6) producers, such as *Pseudomonas bananamidigenes* BW11P2 (36), *Pseudomonas botevensis* COW3 (52) and *Pseudomonas prosekii* LMG26867^T^ (53). Here, the (8:6) refers to the (***l:m***) notation we introduce to provide a simple but effective classification of CLiPs belonging to the same group from a chemical structure perspective, as explained in Materials and Methods. As the stereochemistry of all these bananamides was unknown, we used strain SWRI103 to attempt full structure elucidation, which could only be achieved by the development of our expanded workflow, with all steps in the process described in detail hereafter (Figure S1).

Firstly, analysis of the retrieved BGC predicted a Leu – Asp – Thr – Leu – Leu – Ser – Leu – Ile octapeptide sequence from the associated NRPS (Table 1). The total peptide sequence length and amphipathic profile matches that of previously reported and characterized bananamides (36, 52) but differs in amino acid composition. Therefore, it may represent a novel member of the Bananamide group (8:6). Next, bioinformatic analysis of the condensation/epimerization (C/E) domains was applied for the configurational analysis of the amino acids. The initial C_start_ domain is responsible for the incorporation of a fatty acid (FA) moiety at the N-terminus of the peptide, the exact nature of which cannot be predicted. This domain is followed by six dual activity C/E domains, responsible for the condensation of the newly recruited L-amino acid to the growing peptide chain along with by L to D epimerization of the preceding residue in the sequence (54, 55). This is followed by one ^L^C_L_-type domain, which lacks the epimerization functionality thus retaining the L-configuration of the preceding residue while incorporating an L-amino acid as final residue before cyclisation by a tandem of transesterification (TE) domains. Assuming that all dual C/E domains have functional epimerization activity, the combined A- and C-domain analysis predicts a FA – D-Leu – D-Asp – D-aThr – D-Leu – D-Leu – D-Ser – L-Leu – L-Ile sequence as the most likely lipopeptide biosynthesized by *P. azadiae* SWRI103 (Table 1).

**Table 1:** Sequence, configurational analysis, and assignment of a novel bananamide from *P. azadiae* SWRI103. Underlined amino acids are part of the macrocycle.

| Bananamide from SWRI103 | AA1 | AA2 | AA3 | AA4 | AA5 | AA6 | AA7 | AA8 |
| --- | --- | --- | --- | --- | --- | --- | --- | --- |
| **Bioinformatic analysis workflow** |  |  |  |  |  |  |  |  |
| **A domain** | Leu | Asp | Thr | Leu | Leu | Ser | Leu | Ile |
| **C domain** | C<sub>start</sub> | C/E | C/E | C/E | C/E | C/E | C/E | <sup>L</sup>C<sub>L</sub> |
| **Prediction** | D | D | D | D | D | D | L | L |
| **Chemical analysis workflow** |  |  |  |  |  |  |  |  |
| **NMR analysis** | Leu | Glu | <u>Thr</u> | <u>Leu</u> | <u>Leu</u> | <u>Ser</u> | <u>Leu</u> | <u>Ile</u> |
| **Marfey's analysis** | L/D | D | D | L/D | L/D | D | L/D | L |
| **Combined analysis and synthesized (8:6) Lx library** |  |  |  |  |  |  |  |  |
|  | Leu | Glu | <u>Thr</u> | <u>Leu</u> | <u>Leu</u> | <u>Ser</u> | <u>Leu</u> | <u>Ile</u> |
| **(8:6) L1 (1)\*** | L | D | D-a | D | D | D | L | L |
| **(8:6) L4 (2)\*** | D | D | D-a | L | D | D | L | L |
| **(8:6) L5 (3)\*** | D | D | D-a | D | L | D | L | L |

\* The nomenclature of the synthetic compounds is based on the (*l:m*) notation of the Bananamide group (8:7), followed by the position of the elusive L-Leu in the sequence (position 1, 4 or 5).

Secondly, and independently from the bioinformatic analysis workflow, we engaged into the chemical analysis workflow of the putative novel bananamide. Following incubation of *P. azadiae* SWRI-103 in M9 minimal salt medium, a single CLiP-containing fraction could be extracted and purified. From high resolution MS analysis, a molecular mass of 1053.67 Da could be established i.e., within the expected range for an (8:6) CLiP. Next, NMR was used to elucidate the planar structure. That is, combined COSY/TOCSY analysis showed the presence of a single glutamic acid, serine, threonine, and isoleucine and four leucine residues in the peptide sequence while NOESY and ^1^H-{^13^C} HMBC experiments independently allowed to place these in the same order as predicted from the A-domain analysis, validating the latter (Table 1). Note however that while an aspartic acid residue was predicted at position 2 from the bioinformatic analysis, the presence of a glutamic acid was observed here using NMR spectroscopy. This is not surprising, as it was previously shown that Asp/Glu selectivity from genomic predictions is not always clear cut (36). Additionally, the ^1^H-{^13^C} HMBC spectrum showed a clear ^3^J_CH_ correlation between Thr3 Hβ and Ile8 C’ thereby unambiguously establishing that the ester bond occurs between the hydroxyl side chain of threonine and the isoleucine carboxyl end, thus revealing the six-residue macrocycle expected for a member of the Bananamide group. Taking into account that C_52_H_92_N_8_O_14_ was derived as molecular formula from the HR-MS data, a 3-hydroxydecanoic moiety was inferred and confirmed from the NMR data while its linkage to the N-terminus resulted from characteristic ^3^J_CH_ contacts with Leu1 in the ^1^H-{^13^C} HMBC spectrum. As neither MS nor NMR analysis allows establishing the configuration of the individual amino acid residues, we proceeded to Marfey’s method for the analysis of amino acid configuration. Following total hydrolysis of the CLiP, Marfey’s analysis allowed to unambiguously determine the configuration of all uniquely occurring residues. For the four leucines however, a 2:2 D:L ratio was found, indicating configurational heterogeneity and, as a result, the exact distribution of D and L-Leu residues within the oligopeptide sequence remained hidden. Therefore, at the end of the chemical analysis workflow, a total of six distinct sequences which all satisfy the 2:2 ratio but differ in the distribution of D- and L-leucines should therefore be considered. As a result, the definitive elucidation of the stereochemistry and therefore the chemical structure thus reaches a dead-end.

When information regarding a CLiP sequence is available from both bioinformatic and chemical analysis workflows, the configurational ambiguity is sometimes resolved by proposing that one of the sequences issued from chemical analysis matches the bioinformatic prediction (Figure S1). However, such proposals should always be treated with caution since the epimerization functionality of C/E domains can be inactive in a currently unpredictable manner. Moreover, as aptly illustrated by all cases reported here, the predicted configuration from C-domain analysis may not even match those originating from the chemical analysis workflow. Here, for instance, the bioinformatic analysis proposes a 3:1 D:L ratio for the four leucines rather than the 2:2 found experimentally. Nevertheless, the bioinformatic analysis can still provide guidance to narrow down the number of possible diastereomeric sequences by noting that the ^L^C_L_ domains, which lack the epimerization functionality, will maintain the L-configuration of the amino acid introduced at the preceding position (*vide supra*). Since such ^L^C_L_ domain is present in the final NRPS module of *P. azadiae* SWRI103, the penultimate Leu7 residue will invariably retain its L-configuration upon recruitment. This narrows down the number of possible sequences from six to three: (8:6)-L1 (**1**), (8:6)-L4 (**2**) and (8:6)-L5 (**3**) (Table 1), depending on which leucine position features the remaining L-configuration. In all cases, the 3-hydroxy fatty acid tail is assumed to be (R)-configured, as is generally observed for *Pseudomonas* CLiPs. Notwithstanding the combination of bioinformatic and chemical analysis workflows, the stereochemistry remains ill-defined.

#### 1.2. Extending the chemical analysis workflow by a synthesis add-on

As recently reviewed by Götze and Stallforth, several approaches exist to tackle the issue of configurational ambiguity (24). In the absence of suitable crystals for X-ray diffraction-based structure elucidation, the most common one resorts to mild acid catalyzed hydrolysis conditions or enzymatic degradation of linearized lipopeptides to yield oligopeptide fragments. These then need to be isolated, characterized by NMR or MS, and subjected to Marfey’s method to identify the correct position of the corresponding D- or L-amino acids, introducing an extensive workload without guarantee that the matter will be settled (26). An alternative solution using MS makes clever use of deuterated amino acids and the fact that D-amino acids are, with very few exceptions, generated during biosynthesis through epimerization from L-amino acids (23, 56). In all cases, a separate analysis should be performed to elucidate the stereochemistry of the 3-hydroxy fatty acid moiety (57). Here, we rather build on our previously developed total chemical synthesis route for Viscosin (9:7) group CLiPs (47, 58) to synthesize the three remaining (8:6)-Lx (x=1, 4, 5) sequences (Table 1). The total chemical synthesis strategy mostly relies on solid phase peptide synthesis, affording considerable automation and rapid access to multiple homologous sequences through parallel synthesis (47, 58, 59). With one residue less in the macrocycle and a D-Glu2 rather than D-Gln2 as main differences, the synthesis of the (8:6)-Lx sequences could proceed with minimal change to the original strategy. A more extensive discussion of the applied synthesis route and its key features can be found in the Supplementary Materials section.

Next, the exact stereochemistry of the natural compound is revealed by matching the NMR spectral fingerprint to those of the synthetic compounds. The NMR matching approach relies on previous observations where, using the ^1^H-{^13^C} HSQC experiment, we established that the inversion in the configuration of a single Cα atom introduces prominent changes in the ^1^H and ^13^C chemical shifts of the corresponding (C–H)α group, even when the three-dimensional structure of the CLiP was retained (46). The same trend was found for the stereochemistry of the 3-hydroxy fatty acid moiety (47). Thus, the (C– H)α fingerprint region of the HSQC spectra of the natural compound and its synthetic (8:6)-Lx analogue are expected to be identical. In contrast, the two other sequences will display clearly distinct HSQC spectra in general and (C–H)α fingerprint regions in particular as they feature two D:L inversions compared to the natural compound (Table 1).

By individually overlaying the ^1^H-{^13^C} HSQC spectra of each synthesized (8:6)-Lx variant with that of the natural compound from *P. azadiae* SWRI103, a straightforward visual assessment concerning similarities and differences in the ^1^H and ^13^C chemical shifts can be made (Figure 1). Being the most sensitive reporters of backbone stereochemistry, we focus here on the cross-peaks in the (C-H)α fingerprint regions (Figure 1). Accordingly, the (8:6)-L4 (**2**) variant shows an excellent match with the (C–H)α fingerprint of the natural compound since all (C-H)α cross peak pairs belonging to the same residue in the sequence show excellent overlap (Figure 1A). In contrast, a considerable mismatch exists with (8:6)-L1 (**1**) as visualized by five clearly non-overlapping cross-peaks (Figure 2B). This mismatch becomes even more pronounced for (8:6)-L5 (**3**), where essentially none of the cross-peaks overlap (Figure 1C). Based on this, we can establish the stereochemistry of the *P. azadiae* SWRI103 bananamide as identical to that of the (8:6)-L4 (**2**) sequence (i.e. 3R-OH C10:0 – D-Leu – D-Glu – <u>D-aThr – L-Leu – D-Leu – D-Ser – L-Leu – L-Ile</u>), revealing that the C/E domain in the fifth module of the NRPS system is non-functional for epimerization.

**Figure 1:**
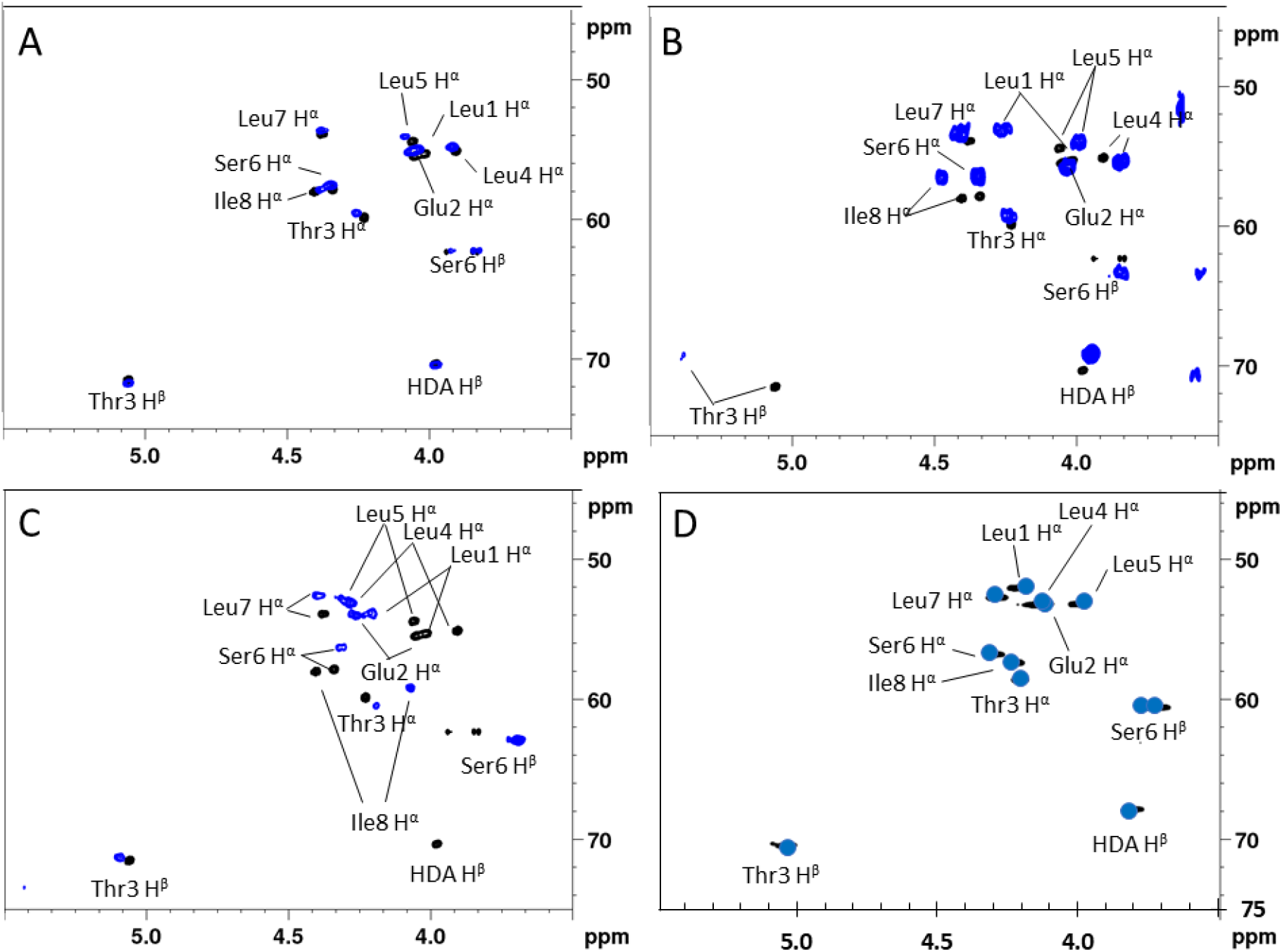
Comparison of the ^1^H-{^13^C} HSQC (CH)α fingerprints of the various synthetic (8:6)-Lx variants (blue) with that of the natural bananamide compound produced by *P. azadiae* SWRI103 (black) recorded in acetonitrile-d3 at 700 MHz, 303K. A) Matching (8:6)- L4 variant B) (8:6)- L1 variant and C) (8:6)- L5 variant. D) shows the spectrum of the natural compound but now recorded in DMSO-d6 at 298K (black) overlayed with a schematic representation of the spectrum of MDN-0066, generated from the tabulated chemical shift data in the original report (17).

**Figure 2:**
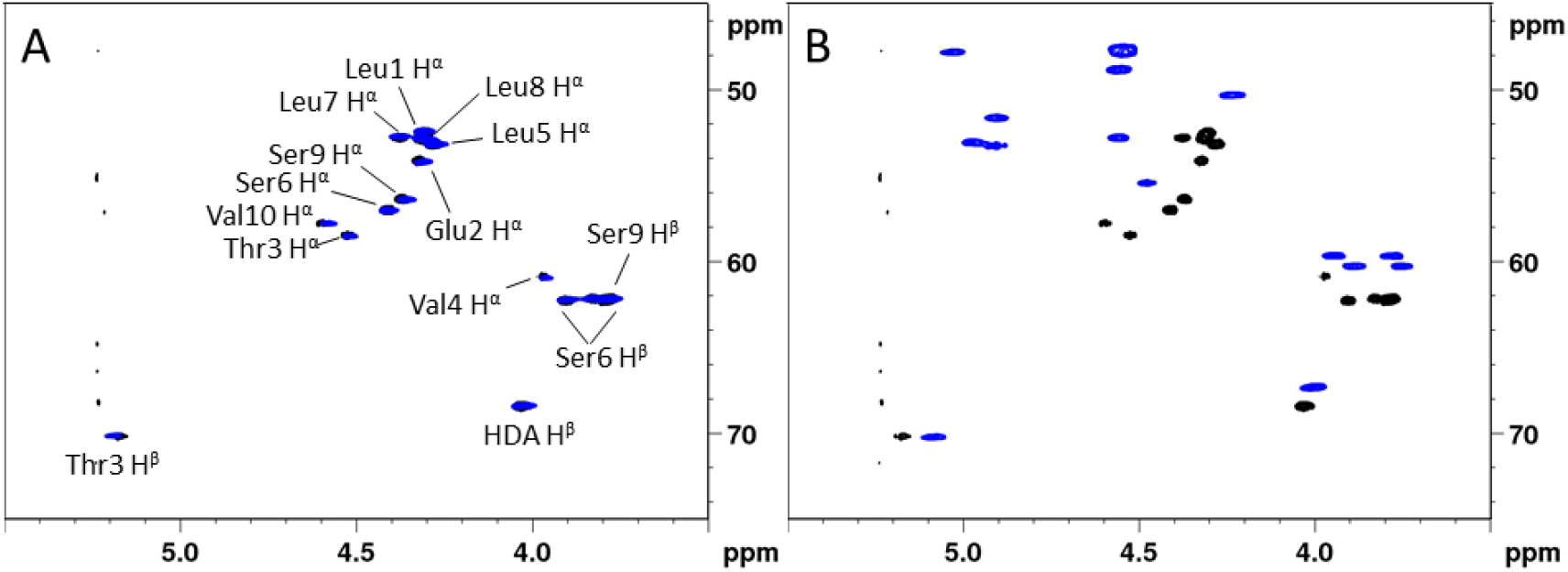
Comparison of the ^1^H-{^13^C} HSQC (CH)α fingerprint of synthetic (10:8)-Lx variants (blue) with that of the natural compound extracted from *P. aestus* CMR5c, all recorded in DMF-d7 at 500 MHz and 298K,. A) Overlay with the synthetic (10:8)-L1 variant (blue) and B) the synthetic (10:8)-L5 variant (blue).

#### 1.3. Matching our reference compound with literature data

The effort to elucidate the stereochemistry of the *P. azadiae* SWRI103 (8:6) bananamide also allows to settle that of MDN-0066, an (8:6) CLiP produced by *Pseudomonas granadensis* F-278,770^T^, which showed distinct bioactivity in a renal carcinoma cell model (17). Using a chemical analysis methodology similar to the one described above, Cautain *et al* elucidated the peptide sequence by relying on MS analysis and NMR spectral assignment. However, the detailed stereochemical elucidation of MDN-0066 was left incomplete as Marfey’s analysis revealed configurational heterogeneity given the presence of 2 D-Leu and 2x L-Leu, a result similar to that of the *P. azadiae* SWRI103 bananamide. Using the tabulated ^1^H and ^13^C NMR chemical shifts reported for MDN-0066 in DMSO-d6, the spectral fingerprint matching could be performed against the data of the SWRI103 bananamide recorded under identical conditions. The result, shown in Figure 1D, shows a quasi-perfect match with (8:6)-L4 (**2**). This proves that MDN-0066 is identical to bananamide SWRI103, thereby disambiguating the stereochemistry of MDN-0066 and illustrating the potential of our approach for dereplication purposes. In order to assess the general applicability of our approach to *Pseudomonas* CLiPs, we then turned to two additional case studies, involving the stereochemical elucidation of orfamide B and xantholysin A.

### 2. Configuration elucidation of (10:8) orfamide B from *P. aestus* CMR5c

Orfamides are important for bacterial motility, cause lysis of oomycete zoospores, and play a role in biocontrol activity against fungal pathogens and insects (6, 48, 60-62). The name-sake of the Orfamide (10:8) group, orfamide A, was extracted from *Pseudomonas protegens* Pf-5 and fully characterized including its complete stereochemistry by mass spectrometry, NMR spectroscopy and chiral gas chromatography. (48) Orfamides B and C were extracted as minors, and their planar structures were characterized as well. Since its original discovery, orfamide B is often found as the major compound produced by newly isolated bacterial strains. (34) Additionally, the orfamide group has expanded with additional orfamide-homologues. (29, 34, 63, 64) However, in many cases, the stereochemistry of orfamide B or that of other orfamide homologues from newly isolated bacterial sources remained unconfirmed as it was derived from sequence similarity with the original orfamide A as extracted from *P. protegens* Pf-5. To unlock conformational analysis and structure-activity evaluations for the Orfamide group, we therefore proceeded to an explicit stereochemical verification.

#### 2.1. Isolation, bioinformatics and initial chemical structure characterization of orfamide B

*P. aestus* CMR5c was originally isolated from the rhizosphere of red cocoyam in Cameroon in a screen for biocontrol agents against the cocoyam root rot disease caused by *Pythium myriotylum (11)*. We already reported the initial genome mining and bioinformatic analysis of the *P. aestus* CMR5c BGC which revealed the presence of three genes with ∼80% similarity to the *ofaA, ofaB* and *ofaC* NRPS genes from *P. protegens* Pf-5 (34). MS and NMR analysis further confirmed the predicted primary sequence and evidenced the incorporation of a 3-hydroxy-tetradecanoic acid moiety at the N-terminus. Thus, the major CLiP produced by *P. aestus* CMR5c has a planar structure identical to the originally published orfamide B. Based on this similarity in primary sequence and the genomic similarity between the respective BGCs, orfamide B from *P. aestus* CMR5c was proposed to also possess only L-leucines (34), as was previously also postulated for the original orfamide A (48).

C-domain analysis showed that the initial C_start_ type domain involved in acylating the first amino acid is followed by six dual activity C/E domains (modules 2–7), two non-epimerizing ^L^C_L_ type domains (modules 8 and 9) and a final C/E domain in the last module (Table 2). Taking into account the distribution of ^L^C_L_ and C/E domains, the bioinformatic analysis predicts the stereochemistry as shown in Table 2, with D-Leu residues occurring at positions 1 and 5 (Table 2) contradicting the all-L-Leu configuration proposed earlier. Marfey’s analysis confirmed the configuration predictions for singly occurring amino acids and the 1:1 D:L ratio for both valines. For the leucines however, a 1:3 D:L ratio was found, invalidating the 2:2 ratio predicted from bioinformatic analysis as well as the all-L configuration proposed by Ma et al (34).

**Table 2:** Sequence, configurational analysis, and assignment of orfamide B from *P. aestus* CMR5c.

| CMR5c | AA<sup>1</sup> | AA<sup>2</sup> | AA<sup>3</sup> | AA<sup>4</sup> | AA<sup>5</sup> | AA<sup>6</sup> | AA<sup>7</sup> | AA<sup>8</sup> | AA<sup>9</sup> | AA<sup>10</sup> |
| --- | --- | --- | --- | --- | --- | --- | --- | --- | --- | --- |
| **Bioinformatic analysis workflow** |  |  |  |  |  |  |  |  |  |  |
| **A domain** | Leu | Glu | aThr | Val/Ile | Leu | Ser | Leu | Leu | Ser | Val |
| **C domain** | Cstart | C/E | C/E | C/E | C/E | C/E | C/E | <sup>L</sup>C<sub>L</sub> | <sup>L</sup>C<sub>L</sub> | C/E |
| **Prediction** | D | D | D | D | D | D | L | L | D | L |
| **Proposal Ma et al. 2016 (33)** | L | D | D | D | L | D | L | L | D | L |
| **Chemical analysis workflow** |  |  |  |  |  |  |  |  |  |  |
| **NMR analysis** | Leu | Glu | <u>Thr</u> | <u>Val</u> | <u>Leu</u> | <u>Ser</u> | <u>Leu</u> | <u>Leu</u> | <u>Ser</u> | <u>Val</u> |
| **Marfey's analysis** | L/D | D | D-a | L/D | L/D | D | L/D | L/D | D | L/D |
| **Synthesized Lx(10:8) library** |  |  |  |  |  |  |  |  |  |  |
|  | Leu | Glu | <u>Thr</u> | <u>Val</u> | <u>Leu</u> | <u>Ser</u> | <u>Leu</u> | <u>Leu</u> | <u>Ser</u> | <u>Val</u> |
| **(10:8)-L1 (4)** | L | D | D-a | D | **D** | D | L | L | D | L |
| **(10:8)-L5 (5)** | **D** | D | D-a | D | L | D | L | L | D | L |
| **(10:8) L1L5 (6)** | L | D | D-a | D | L | D | L | L | D | L |

\* The nomenclature of the synthetic compounds is based on the (*l:m*) notation of the Orfamide group (10:8), followed by the position of the elusive L-Leu in the sequence (position 4, 5 or 4&5).

#### 2.2. Stereochemical elucidation using chemical synthesis and NMR fingerprint analysis

Considering the experimental evidence from the chemical analysis workflow, a total of 8 sequences should effectively be considered since the configurational D:L heterogeneity of the Leu and Val residues are combinatorically independent. Conveniently, the bioinformatic analysis allows to trim this down to two sequences only, strongly reducing the synthetic effort required. Indeed, the ambiguity regarding the valines can be settled by noting that, as the final residue, Val10 is not subjected to any epimerization activity. This pins the valine configurations down as D-Val4 end L-Val10. Next, the presence of ^L^C_L_ domains in module 8 and 9 allows to unequivocally attribute the L-configuration to Leu7 and L-Leu8. The remaining D and L configured leucines are to be distributed over positions 1 and 5. While this constitutes an apparent dead-end, configurational assignment could be finalized using fingerprint matching of the natural compound against the (10:8)-L1 (**4)** and (10:8)-L5 (**5**) sequences obtained by synthesis (Table 2) using the same strategy as discussed above for the bananamide analogues. This included the incorporation of a 3R-hydroxydecanoic moiety (3-OH C10:0) at the N-terminus rather than a 3R-hydroxytetradecanoic one (3-OH C14:0) mainly because of synthetic availability of the precursor. Previous investigation of C_10_, C_12_ and C_14_ pseudodesmin analogues evidenced that lengthening of the acyl chain did not affect the ^1^H and ^13^C chemical shifts of the peptide moiety in any way (59).

Figure 2 shows the overlay of the (C–H) α fingerprint region of each synthesized (10:8)-Lx sequence with that of natural orfamide B from *P. aestus* CMR5c. A straightforward visual assessment clearly indicates that the (10:8)-L1 (**4**) variant displays a quasi-perfect fingerprint match while none of the (C–H)α cross peaks of (10:8)-L5 (**5**) overlap with those of the CMR5c orfamide. The latter probably indicates a major conformational effect caused by the L to D inversion at Leu5. To remove any doubts and independently exclude the all-L-Leu configuration originally proposed from sequence similarity with orfamide B from *P. protegens* Pf-5, we also committed to synthesize the corresponding (10:8)-L1L5 (**6**) analogue which again showed significant mismatch with the natural CMR5c orfamide. (Supplementary figure S36) All data together establish that the stereochemistry of orfamide B from *P. aestus* CMR5c corresponds to that of the (10:8)-L5 (**5**) sequence (3R-OH C14:0 – L-Leu – D-Glu – <u>D-aThr – D-Val – D-Leu – D-Ser – L-Leu – L-Leu – D-Ser – L-Val</u>), indicating the C/E domain in the second module is non-functional for epimerization.

#### 2.3. Stereochemical reassessment and dereplication of the Orfamide group using literature NMR data

The results presented above appear to indicate that orfamide B from *P. aestus* CMR5c is a D-Leu5 diastereoisomer of orfamide B from *P. protegens* Pf-5 (48), since the latter is reported to possess a L-Leu5. So far, reports of CLiPs from the same (***l:m***) group that feature a configurational difference are few, possibly due to the lack of extensive configurational assignment among *Pseudomonas* CLiPs. The best-known example occurs within the Viscosin (9:7) group where CLiPs can be divided in a L-subgroup (14 sequences) and D-subgroup (5 sequences) depending on the configuration of the leucine, also at position 5 (65). Given that orfamides have been reported from multiple bacterial sources since their initial discovery, the question raised as to whether such division is also present for the Orfamide (10:8) group. Settling this matter would allow assessing whether stereochemical variation within a group is a more general feature present in the NRPS of *Pseudomonas* CLiPs. Conveniently, the major orfamides reported from other bacterial sources in literature feature planar structures identical to either orfamide A from *P. protegens* Pf-5 or orfamide B from *P. aestus* CMR5c, thus allowing the use of NMR fingerprinting for stereochemical evaluation. To be able to screen directly against a reference spectrum of orfamide A under standardized conditions, *P. protegens* Pf-5 was cultured to obtain orfamide A, together with a series of minor compounds including orfamide B and 4 previously unreported ones, which we named orfamides J – M (See supplementary information).

For the orfamide B producer *Pseudomonas sessilinigenes* CMR12a (66), the ^1^H-{^13^C} HSQC spectral fingerprint proved identical to the spectrum of the orfamide B reference from *P. aestus* CMR5c. This indicated that these molecules have identical stereochemistry and feature a D-Leu5. Next, we turn to *Pseudomonas* sp. PH1b where genome exploration revealed an orfamide-type BGC (67). Characterization of orfamides from this isolate presents an interesting case as it produces a major and a minor orfamide with sequence identical to orfamide A and orfamide B, respectively, thus only differing by a Ile4/Val4 substitution resulting from A-domain substrate flexibility in module 4 of the NRPS. The (C–H)α spectral fingerprint of its major orfamide matched with that of orfamide A produced by *P. protegens* Pf-5 indicating an L-configuration for Leu5. Surprisingly, however, the spectrum of the minor orfamide of *Pseudomonas* sp. PH1b matched with that of orfamide B from *P. aestus* CMR5c, therefore indicating a D-Leu5 configuration. This result was highly unexpected as it implies that in *Pseudomonas* sp. PH1b, the same NRPS assembly line would yield different configurations for Leu5. More specifically, it would require the occurrence of epimerization by the C/E domain of module 6 to correlate with the sequence composition of the growing peptide chain as recruited by module 4. Since the all-L Leu configuration of the original orfamide A from *P. protegens* Pf-5 was derived from extensive chemical analysis but not confirmed through total synthesis, it appeared more likely that an error was made in establishing the configuration at this position. To settle this matter, we reinvestigated the stereochemistry of the original orfamide A, as extracted from *P. protegens* Pf-5. In contradiction with the report by Gross *et al*, where chiral GC revealed a 0:4 D:L ratio for the leucines, our Marfey’s analysis yielded a 1:3 D:L ratio, definitively settling the case in favor of a D-Leu5. As a result, in this case, configurational variability within one NRPS assembly line can be invalidated, and a revision is required for the configuration of Pf-5 based orfamides as shown in Table

Gratifyingly, this outcome was recently also arrived at using a completely independent approach involving total synthesis of orfamide A by Bando et al. (*personal communication*), whereby the original stereochemistry led to loss of green algal deflagellation activity, while the corrected one was as active as the natural compound (68). Finally, we could establish that the orfamide produced by *Pseudomonas* sp. F6 (8) is identical to the revised orfamide A, by only making use of the tabulated chemical shift values of the published compound, and matching this to our reference spectrum of orfamide A. (Supplementary figure S41)

### 3. Settling the elusive stereochemistry of xantholysin A

An additional example illustrating the strength of our combined synthesis and spectral matching approach concerns xantholysin A. First reported as the main CLiP produced by *Pseudomonas mosselii* BW11M1, together with minor congeners B to D, it is the prototype that defines the Xantholysin (14:8) group (69). Since no configurational analysis was performed at the time, NMR and MS analysis only provided the planar structure. Subsequent reports proposed the production of xantholysins by other *Pseudomonas* strains, uncovering a diverse portfolio of biological functions in the process. Pascual *et al* (16) reported the production of xantholysins A to D by *Pseudomonas soli* F-279,208^T^ solely based on HR-MS of the isolates and characterized their cytotoxic activity against the RCC4 kidney carcinoma cell line. Lim *et al* (9) reported HR-MS and ^1^H NMR identical to those of xantholysin A and B for two lipopeptides from *Pseudomonas* sp. DJ15 and demonstrated their insecticidal activity against the green peach aphid, a major peach tree pest also serving as plant virus vector. In the context of exploring the tolerance of cocoyam plants against *Pythium myriotylum* in the field, Oni *et al*. (43) showed that *Pseudomonas* sp. COR51 produces a lipopeptide with identical planar structure to that of xantholysin A. It was also extracted and characterized from *Pseudomonas xantholysinigenes* in the framework of the development of a diagnostic bioinformatics tool that allows the assignment of a CLiP to a particular lipopeptide group based on the phylogeny of the *MacB* transporter of its producing bacterium. (53) In other work, a large-scale bacterial screening for antibiotic activity proposed the identification of xantholysins A-D in *Pseudomonas* sp. 250J, based on HR-MS data and high similarity in the NRPS genomic make-up to that of *P. mosselii* BW11M1 (70). Later on, combination of Marfey’s analysis with mapping of the C/E and ^L^C_L_ domains in its NRPS led to propose a stereochemistry for xantholysin A, although the distribution of the remaining D-Leu and L-Leu over positions 9 and 11 remained tentative (71). Shimura *et al* (72) reported the total synthesis of MA026, a xantholysin-like CLiP from *Pseudomonas* sp. RtlB026 with potent anti-hepatitis C activity. Discrepancies in physicochemical properties of synthetic and natural MA026 led Uchiyama *et al*. (73) to revise its structure by exchanging residues at position 10 and 11, as previously suggested by Li *et al* (69), thus leading to a planar structure identical to xantholysin A. With more than 90% sequence similarity between their respective BGC, it was proposed that the stereochemistry of xantholysin A would be identical to that of MA026. However, Uchiyama *et al*. also noted that the stereochemical assignment for xantholysin A from *Pseudomonas* sp. 250J did not match with MA026, the latter containing L-Gln6 and D-Gln13, while the reverse configuration was proposed for the former.

To resolve this on-going characterization issue, we first considered applying our combined approach to the original xantholysin A from *P. mosselii* BW11M1. However, the presence of configurational heterogeneity for the leucine (D:L 2:1) and glutamine/glutamic acid (Glx) residues (D:L 4:1) independently act to generate 50 possible sequences, that can be trimmed down to 15 sequences when taking into account the genomic data. However, no further prioritization is possible after combining the chemical and genomic data. Comparison of the (C–H)α fingerprint of MA026 (with known stereochemistry) to that of originally isolated xantholysin A should in principle allow to either quickly establish configurational identity or eliminate one sequence. However, NMR data for MA026 was listed without explicit assignment. To circumvent this, we adapted our own synthesis scheme to produce MA026 and subsequently established that its (C–H)α fingerprint is indeed identical with that of the original xantholysin A (Figure 3), thus avoiding the need for further synthesis. Used as reference compound, simple comparison of its (C–H)α fingerprint with that of putative xantholysin A lipopeptides allowed to also extend the assigned stereochemistry to the major xantholysin extracted from *Pseudomonas*. sp. COR51 (43), *P. xantholysinigenes* (53) and *Pseudomonas* sp. 250J (71), for which the NMR data were available or provided for comparison by the original authors, respectively. (See Supplementary figures S57-S59) In conclusion, the structure of xantholysin A was determined to be 3R-OH C10:0 – L-Leu – D-Glu – D-Gln – D-Val – D-Leu – L-Gln – <u>D-Ser – D-Val – D-Leu – D-Gln – L-Leu – L-Leu – D-Gln – L-Ile</u>. This reveals that two modules (2 in *XtlA* and 7 in *XltB*) lack the epimerization capability expected for the respective C/E-classified domains.

**Figure 3:**
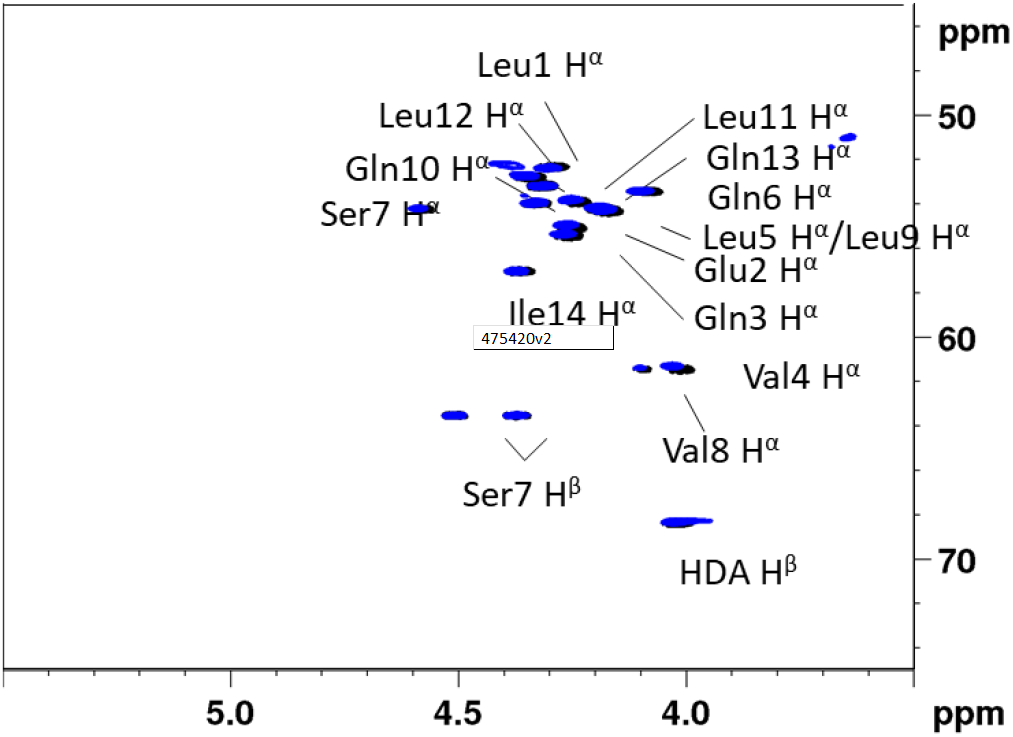
Comparison of the ^1^H-{^13^C} HSQC (C-H)α fingerprints of natural xantholysin A (14:8) produced by *P. mosselii* BW11M1 (black) with that of MA026 obtained through synthesis (blue). DMF-d7, 328K, 700MHz.

## Discussion

The discovery and development of new antimicrobial agents and therapies is an important avenue to tackle the rising global health threat caused by antimicrobial resistance (74, 75). Because of their broad range of antimicrobial activities and wide diversity of structures, *Pseudomonas* CLiPs have drawn attention as an interesting class of compounds for further exploration in this respect (34, 59, 76, 77). To engage into structure-activity-relationship (SAR) studies, complete elucidation of their structure, including stereochemical make-up, is of vital importance to identify meaningful structure-activity associations (34, 59) and uncover their mode of action. Unfortunately, incomplete structure characterization increasingly accompanies the rising number of these CLiPs isolated from novel bacterial sources. Mass spectrometric (MS) data, genomic predictions or a combination thereof yield insufficient structural data by itself to generate a single structure elucidation result. Especially the comparative analysis with pre-existing data to establish structural novelty or identity with existing CLiPs is not approached critically enough to avoid erroneous attributions. Consequently, discovery efforts may generate a narrowed view on lipopeptide structural diversity due to erroneous assignment of *de facto* new structures to previously described ones. In turn, this clouds the comparative analysis and interpretation of antimicrobial activities of newly extracted compounds with already existing ones and threatens to lead subsequent development efforts astray. As natural product mining efforts aimed at *Pseudomonas* strains amplify, the chance of rediscovery of existing CLiPs increases as well. As a result, complete characterization including the labor-intensive determination of stereochemistry – while essential – may not provide a ‘return-on-investment’ as much effort might be spent on an already characterized compound. (Figure 4A)

**Figure 4:**
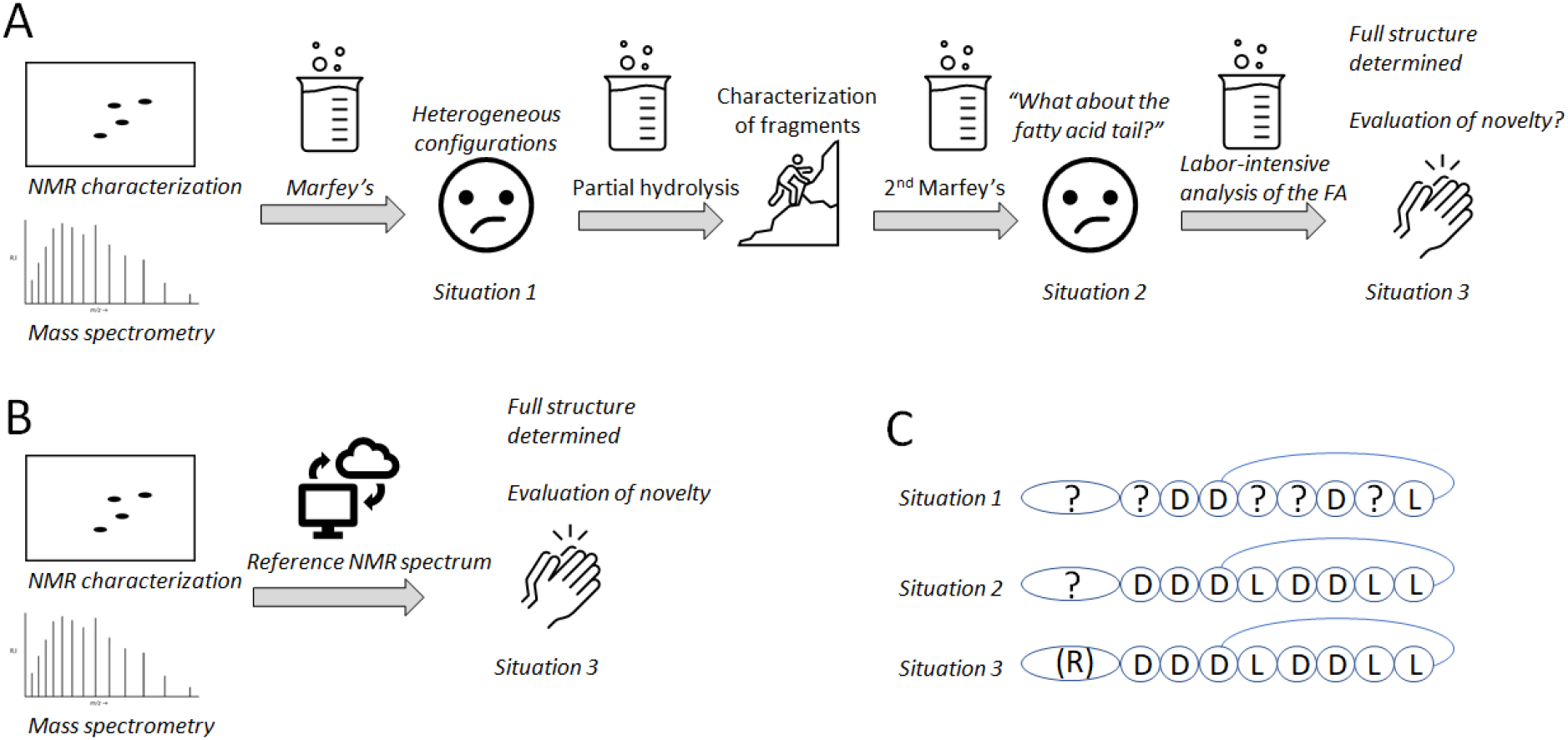
The stereochemical analysis of CLiPs requires a series of analysis steps. A) Chemical analysis steps typically required before the implementation of our NMR fingerprint matching methodology described here. B) Using NMR fingerprint matching, a single analysis step is required for identity confirmation as well as stereochemical validation. C) Different levels of stereochemical validation of CLiPs.

As part of a concerted research effort aiming to map the structural and conformational diversity of *Pseudomonas* CLiPs, we extended the existing characterization workflows by a combined synthesis and NMR fingerprint matching approach. Simultaneously, it allowed us to demonstrate and validate the innate dereplication potential of the latter by a simple screening of the NMR fingerprint of newly isolated CLiPs. First, we established that the CLiP extracted from the newly isolated producer *P. azadiae* SWRI103 belongs to the Bananamide (8:6) group and subsequently resolved a configurational dead-end to reveal its stereochemistry. By utilizing our NMR spectral matching methodology, it was subsequently found to be identical to MDN-0066 produced by the *P. granadensis* type-strain, thereby settling the stereochemistry of this compound as well (17). Using the same approach, we determined that a structure revision is required for several, if not all, Orfamide (10:8) group members. While initially only poised to establish the stereochemical make-up of orfamide B as extracted from *P. aestus* CMR5c on a firmer basis, we showed that this compound possesses a D-Leu5, instead of an L-Leu5 as previously presumed and published (34, 48). Comparing the ^1^H-{^13^C} HSQC spectral fingerprint of a collection of CLiPs produced by *P. protegens* Pf-5, *P. sessilinigenes* CMR12a, *Pseudomonas* sp. F6 and *Pseudomonas*. sp. PH1b that share the orfamide sequence but have unreported stereochemistry, the presence of a D-Leu5 was confirmed in all cases, proving that one and the same compound is produced by diverse strains from a variety of environments. Given an L-Leu5 was now confined to the originally reported orfamide A only, NMR fingerprint matching allowed to convincingly assert that here also, a D-Leu5 should be considered. Moreover, it was determined that all orfamide members possess a 3R-hydroxy-fatty acid tail, in contrast to earlier literature reports asserting the presence of a 3S-hydroxy-fatty acid tail (48, 57). Finally, we determined the stereochemistry of xantholysin A, as produced by a variety of sources. Notably, with 15 possible sequences resulting from the combination of information obtained from both genomic as well as chemical analysis protocols, a considerable synthetic effort would have been required for the elucidation of this lipopeptides’ stereochemistry. Instead, we utilized available literature data to prioritize possible sequences, resulting in only a single preferred sequence to be initially synthesized, thereby solving the stereochemistry of xantholysin A (as produced by various bacterial strains; using their reported literature data) and MA026.

From the above, it is clear that the NMR fingerprint approach introduced in this work will support researchers in dereplication i.e., has this particular lipopeptide been isolated before? By matching NMR spectra of a CLiP from a newly isolated bacterial source with those of existing (reference) CLiPs, one can determine whether they are identical or not. Importantly, this is possible irrespective of the stereochemical characterization of the reference CLiPs and does not require any synthesis or extensive characterization effort. In fact, it can be performed at a relatively early stage following isolation and purification, as it only requires a ^1^H-^13^C fingerprint, which is preferably but not necessarily assigned. Indeed, in case of a perfect match, identity is ensured, while small but notable differences may motivate further characterization using the workflow as described here (Figure S1)

It is important to note that total synthesis is not a prerequisite for dereplication of a newly isolated CLiP. In other words, the (C–H) α NMR fingerprint matching only requires the comparison of NMR spectra, or tabulated ^1^H and ^13^C chemical shifts values to assess structural similarity to a reference CLiP which may arise from natural sources or be synthetic compounds issued from structure elucidation efforts such as described in this work. Provided the required data is collected and made accessible, a fast screening method becomes available early on in the discovery process, potentially eliminating additional labor-intensive analyses, especially when it concerns CLiPs with a configurational heterogeneity of the amino acid content. Only when a genuinely new CLiP structure is found and/or no reference compound is available, additional analysis is desired/required. In such cases, we have shown that genomic and chemical methods, in combination with solid-phase peptide synthesis can provide unambiguous characterization of CLiP structures.

Application of NMR based dereplication as introduced here should permit an improved mining of NRPSs by providing reference data for bioinformatics software suites, thereby improving genomic prediction of CLiP structures. As previously observed for *Pseudomonas* lipopeptides from the Viscosin, Poaeamide, Tolaasin, Factin, Peptin, and Mycin families (78, 79) and further evidenced in this report for CLiPs from three additional groups (Bananamide, Orfamide, Xantholysin), the bioinformatic prediction of epimerization functionality in the non-ribosomal biosynthesis of CLiPs remains inaccurate. In each of these groups, at least one module appears catalytically inactive for epimerization. Only in the Amphisin (11:9) and Gacamide/Cocoyamide (11:5) groups, do the predictions of dual activity condensation domains match with the elucidated CLiP stereochemistry. It can be anticipated that ensuring availability of unambiguous stereochemical information and scrutinizing corresponding C/E domain sequences for motifs linked to epimerization (in)activity (22), ideally complemented with insight from currently unavailable 3D structural C/E domain data, should allow an improved genomic prediction of the affected amino acid configurations.

## Conclusion

Incomplete (or erroneous) structural characterization currently generates ‘scientific noise’ in the field of lipopeptide research, hampering a concerted research action for the generation of structure-activity relationships (18). The (C–H)α NMR fingerprint matching approach as described here will prevent this by offering a rapid, unambiguous characterization of CLiP structures. By standardizing the relevant experimental conditions as much as possible and by making reference data publicly available, a rapid elucidation of the complete structure of newly extracted lipopeptides, including stereochemical make-up of both the oligopeptide and the fatty acid moieties comes within reach. (Figure 4) In addition, our user friendly NMR matching methodology allowed to quickly supplement incomplete literature and characterization data with NMR fingerprint data, thereby elucidating the complete structure (including stereochemistry) of previously extracted CLiPs. (Figure S2) While limited to *Pseudomonas* CLiPs, our results support the (C– H) α NMR fingerprint matching as a tool for dereplication beyond this genus, such as those produced by other bacterial phyla, including *Bacillus, Burkholderia* and *Streptomyces* spp. Obviously, access to the desired NMR data for a sufficiently large collection of CLiPs, preferably with annotated fingerprints, is essential for this approach to become of use for the research community. Only then can the lack of a match be interpreted as indicative of compound novelty with high confidence. While awaiting the development of a (dedicated) NMR data sharing platform – most likely by integration of the reference spectra into existing (lipopeptide)databases, such as Norine (80, 81) – we have made a publicly available ‘knowledge base’ at https://www.rhizoclip.be, where we present an overview of downloadable (C–H)α NMR fingerprint data of characterized CLiPs, together with literature data on the originally determined structures. The latter includes a description of the CLiPs’ original description, molecular mass, three-dimensional structures (if available), and a summary of published antimicrobial activities. Moreover, a detailed protocol is available for researchers that wish to record NMR data of their newly extracted lipopeptides to compare them to the publicly available reference data. Finally, we invite all researchers to submit NMR data of new CLiPs, such that they can be included in this knowledge base.

## Materials and Methods

Detailed materials and experimental methods can be found in the SI appendix together with additional tables and figures.

## Data Availability

All study data are included in the article and *SI Appendix*. NMR data for (C–H)α spectral fingerprint matching can be found at https://www.rhizoclip.be

## Acknowledgments

AM and JM are recipient of a concerted research action grant (MEMCLiP) from Ghent University supporting this research (GOA-028-19). JM and MH are recipients of the Excellence of Science grant RhizoCLiP (EOS ID 30650620) supporting this research. The NMR equipment used in this work was funded through the FFEU-ZWAP initiative and grants from the Hercules Foundation (AUGE09/06 and AUGE15/12) awarded to a consortium headed by J. Martins. The NMR equipment is part of the NMR Centre of Expertise at UGent. We would like to thank Pauline De Coster and Ilse Delaere for practical assistance with bacterial cultures, HPLC purifications and NMR characterization. Thanks are extended to Kok-Gan Chan (University of Malaya, Malaysia) for use of the *Pseudomonas* sp. PH1B strain and Carlos Molina Santiago for providing the NMR spectral data of xantholysin A as extracted from *Pseudomonas* sp. 250J for fingerprint matching purposes.

## Notes

https://www.rhizoclip.be

### Competing Interest Statement

The authors have declared no competing interest.

